# CsrA-controlled proteins reveal new dimensions of *Acinetobacter baumannii* desiccation tolerance

**DOI:** 10.1101/2021.08.11.455981

**Authors:** Yasuhiro Oda, Madelyn M. Shapiro, Nathan M. Lewis, Xuefei Zhong, Holly K. Huse, Weizhi Zhong, James E. Bruce, Colin Manoil, Caroline S. Harwood

**Affiliations:** Department of Microbiology, University of Washington, Seattle, Washington, 98195-7735 USA; Seattle Children’s Research Institute, Seattle, Washington, 98109 USA; Department of Plant and Microbial Biology. University of Minnesota, St. Paul, MN. 55108 USA; Analytical Chemistry Group. Regeneron Pharmaceuticals, Tarrytown, NY 10591 USA; Department of Pathology. Harbor-UCLA Medical Center, Torrance, CA 90502-2059 USA; Department of Genome Sciences, University of Washington, Seattle, Washington, 98195-5065 USA

**Keywords:** *Acinetobacter baumannii*, desiccation, CsrA, intrinsically disordered proteins

## Abstract

Hospital environments are excellent reservoirs for the opportunistic pathogen *Acinetobacter baumannii* in part because it is exceptionally tolerant to desiccation. We found that relative to other *A. baumannii* strains, the virulent strain AB5075 was strikingly desiccation resistant at 2% relative humidity (RH), suggesting that it’s a good model for studies of the functional basis of this trait. Consistent with results from other *A. baumannii* strains at 30% RH, we found the global post-transcriptional regulator CsrA to be critically important for desiccation tolerance of AB5075 at 2% RH. To identify CsrA-controlled proteins that may contribute to desiccation tolerance we used proteomics to identify proteins that were differentially present in wild type and *csrA* mutant cells. Subsequent mutant analysis revealed nine genes that were required for wild type levels of desiccation tolerance, five of which had modest phenotypes. Catalase and a universal stress protein gene were moderately important for desiccation tolerance and two genes of unknown function had very strong desiccation phenotypes. The predicted amino acid sequence of one of these genes predicts an intrinsically disordered protein. This category of proteins is widespread in eukaryotes but less so in prokaryotes. Our results suggest there may be mechanisms responsible for desiccation tolerance that have not previously been explored in bacteria.

**IMPORTANCE:** *Acinetobacter baumannii* is commonly found in terrestrial environments but can cause nosocomial infections in very sick patients. A factor that contributes to the prevalence of *A. baumannii* in hospital settings is that it is intrinsically resistant to dry conditions. Here, we established the virulent strain *A. baumannii* AB5075 as a model for studies of desiccation tolerance at very low relative humidity. Our results show that this trait depends on two proteins of unknown function, one of which is predicted to be an intrinsically disordered protein. This category of protein is critical for the small animals named tardigrades to survive desiccation. Our results suggest that *A. baumannii* may have novel strategies to survive desiccation that have not previously been seen in bacteria.

## INTRODUCTION

Hospital-acquired infections are an important healthcare concern and economic burden (1, 2) and environmental persistence plays a critical role in the transmission of bacteria that cause these infections (3–6). One such bacterium is *Acinetobacter baumannii*, an opportunistic pathogen that infects very sick patients. It is responsible for about 2% of nosocomial infections in the United States and Europe and the frequencies are higher in the rest of the world. *A. baumannii* is especially problematic because on a global basis, about 45% of isolates are multi-drug resistant (7). A factor that contributes to the prevalence of *A. baumannii* in hospital settings is desiccation tolerance. *A. baumannii* can survive in a desiccated state on inanimate dry surfaces for days to several months (8–10). These surfaces include materials that are often encountered in the hospital, such as polyvinyl chloride, rubber, and stainless steel (11).

When desiccated, bacteria must respond to diverse stresses that include accumulation of reactive oxygen species, loss of cytoplasmic volume, and loss of cell membrane integrity (12, 13). Proteomics analysis of *A. baumannii* showed that desiccated cells had higher levels of proteins involved in protein stabilization, antimicrobial resistance, and reactive oxygen species detoxification (14). Attributes of *A. baumannii*, that have been shown to be associated with desiccation tolerance include biofilm formation (15, 16) and protein aggregation (17). LpxM_AB_-dependent acetylation of lipid A is essential for survival of *A. baumannii* ATCC17978 at 40% RH (18), and a *recA* mutant of ATCC17978, defective in DNA repair, had a pleiotropic phenotype, including a defect in desiccation tolerance (19). *katE*, encoding catalase also contributes to desiccation tolerance (20). Despite these observations, the number of genes identified in *A. baumannii* that are specifically involved in desiccation tolerance is small. This could be because *A. baumannii* cells have evolved modified cell structures that are both essential for viability and important for desiccation tolerance. The genetic basis for this would be difficult to uncover in mutant screens. It is also possible that the conditions of desiccation used, typically 30% RH in studies to date, were not sufficiently severe to allow identification of some desiccation tolerance genes.

With the goal of identifying new desiccation tolerance genes, we established *A. baumannii* strain AB5075 as a model for studying desiccation tolerance under severe conditions of 2% RH. We then followed up on a recent report showing that *csrA*, which encodes a global post-transcriptional regulator, is important for desiccation tolerance (21) and identified nine CsrA-controlled proteins that confer desiccation tolerance on AB5075 at 2% RH. One of these has predicted properties that suggest new dimensions of desiccation tolerance.

## RESULTS

### Desiccation assay

Previous studies have shown that *A. baumannii* can survive in a desiccated state for days to several months (8–11, 20). For these and other desiccation studies, investigators worked with a variety of strains and usually incubated cells at either 30% RH or in room air, which varied between 25 and 61% RH in one study (20). These differences can make it difficult to compare desiccation phenotypes between studies. Thus we thought it important to establish a robust desiccation assay that reduces experimental variables like choice of strains, drying times, and RH during desiccation, using a highly desiccation resistant strain.

Following from previous reports, saturated calcium chloride hexahydrate solution placed in a sealed plastic Snapware container caused the RH inside the container to rapidly equilibrate to 30% (9). We found that use of DRIERITE instead of calcium chloride, resulted in an RH of 2%. To test desiccation tolerance under different conditions, we grew bacteria to a desired density in Trypticase Soy Broth (TSB), harvested them, washed them twice, and resuspended them in phosphate buffer to a final OD_600_ of 1. Drops of cell suspension were placed on polycarbonate membranes and filtered to allow for rapid drying. The membranes were then placed in uncapped 15 ml conical centrifuge tubes and incubated in desiccation containers. After various periods of incubation, buffer was added to each centrifuge tube followed by 5 min of shaking on a rotary shaker. Viable cell numbers were then determined by plating on TSB agar. To control for the stress of filtration we did viable counts immediately following filtration and took this as our “day 0” time point.

### Relative desiccation tolerance of *A. baumannii* strains

As shown in Fig 1A, *A. baumannii* strain AB5075 and *Escherichia coli* strain W3100 each survived desiccation at 30% RH far better than *Pseudomonas aeruginosa* strain PAO1. However at 2% RH, *A. baumannii* survived far better than either *E. coli* or *P. aeruginosa*. As has been reported (20, 22), we found that *A. baumannii* stationary phase cells were much more tolerant to desiccation than actively growing cells (Fig. S1) and so we routinely used stationary phase cells in our desiccation assays. We compared the desiccaton tolerance of AB5075 to two additional frequently used laboratory strains of *A. baumannii*, ATCC17978 and ATCC19606. We found that AB5075 was strikingly more resistant than the others at 2% RH and somewhat mores resistant at 30% (Fig. 1B). Given the strength of its phenotype, these findings indicate that AB5075 is a good model for studing desiccation tolerance. This strain was isolated from a surgical wound, is multidrug resistant and is highly virulent in an animal model (23). A comprehensive ordered mutant library of AB5075 is available that has two to three seqenced Tn insertions in each gene and is called the three-allelle library (24).

**Figure 1.**
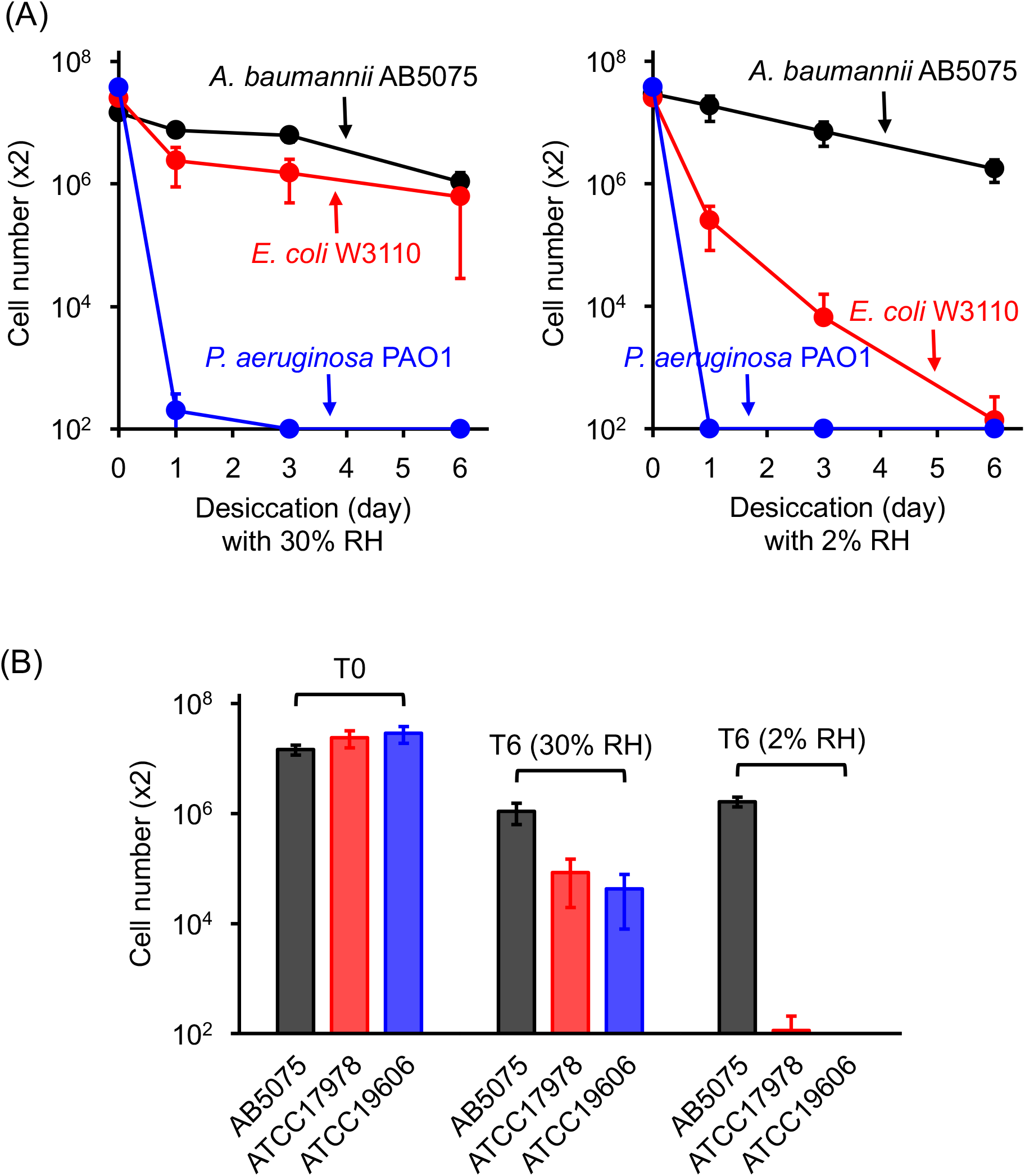
Desiccation tolerance of *A. baumannii*. (A) Comparison of AB5075, *E. coli* W3110, and *P. aeruginosa* PAO1 at 30% RH or 2% RH. (B) Desiccation tolerance of *A. baumannii* strains AB5075, ATCC17978, and ATCC19606 after 0 days (control) and 6 days of desiccation at 30% RH or 2% RH. The cell numbers represent the total number of viable cells recovered from each membrane. The data are the average of three or more biological replicates, and standard deviations are shown as error bars.

### CsrA is critical for desiccation tolerance

We examined the contribution of the post-transcriptional regulator CsrA to desiccation tolerance of AB5075 by constructing a *csrA* deletion mutant (Δ*csrA*). We found that the mutant grew poorly on TY agar and had an elongated cell morphology when grown in TY broth (Fig. 2A). On agar plates, large colonies frequently appeared on a backgound of poor growth, likely due to occurance of second site suppressor mutations in the Δ*csrA* strain. The Δ*csrA* strain was also defective in growth on other nutrient-rich media, including LB broth, nutrient broth, and TSB broth. A similar sensitivity to growth in complex media was reported by Farrow et al. for several *A. baumannii* strains including strain AB5075 (21). In agreement with Farrow et al., a Δ*csrA* mutant grew as the wild type in defined medium, in our case, M9 minimal medium with 10 mM succinate as a sole carbon source (M9/succinate), and it had close to a wild type cell morphology (Fig 2A). A *Yersinia enterocolitica csrA* mutant, has a growth defect in LB due to the presence of 90 mM of NaCl (25). However, the *A. baumannii* Δ*csrA* mutant was not sensitive to this level of NaCl. In fact, the mutant grew in M9/succinate supplemented with up to 100 mM of NaCl without a significant reduction of growth compared to the wild type.

**Figure 2.**
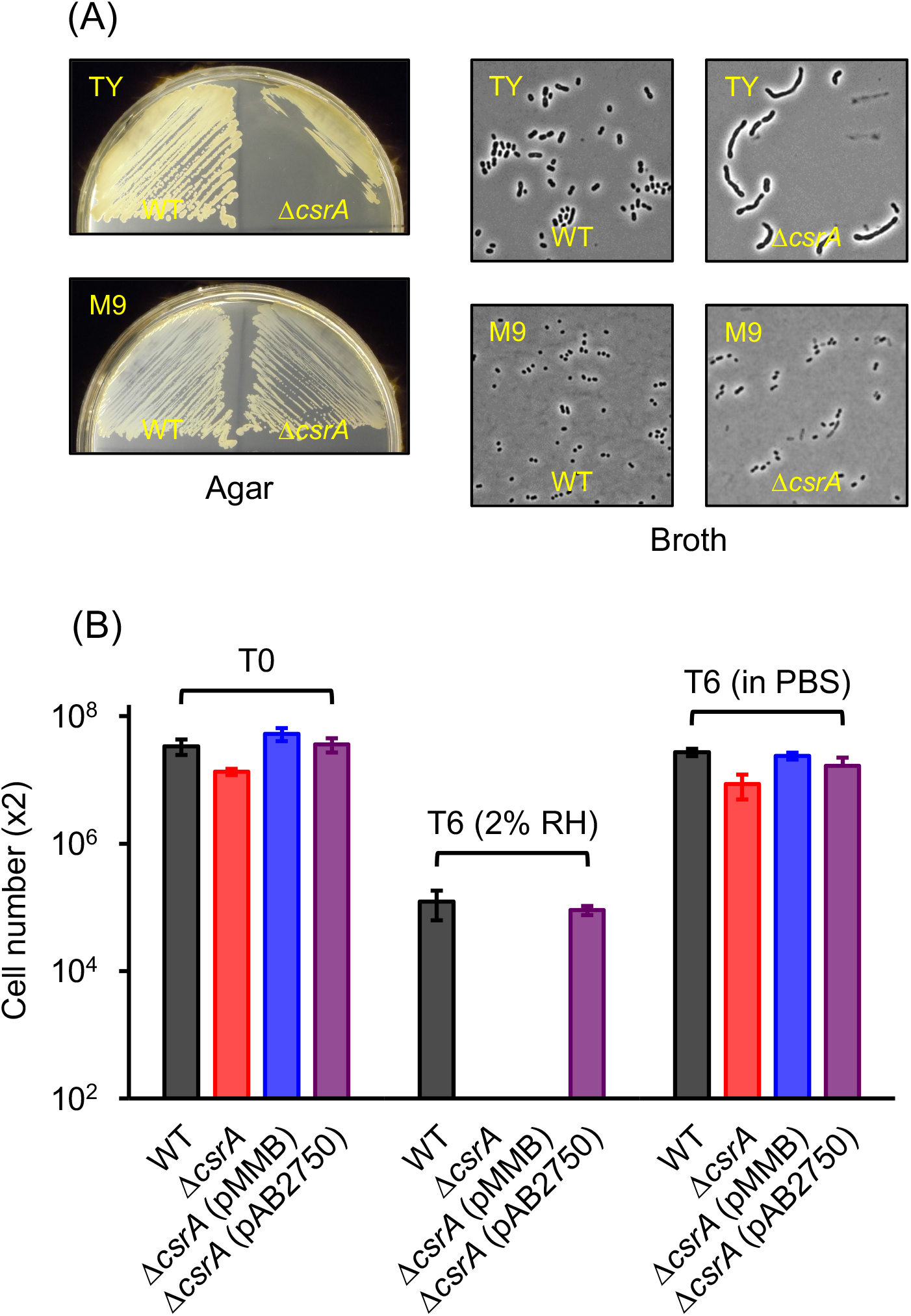
Growth and desiccation tolerance of a Δ*csrA* mutant (A) Comparison of wild type (WT) and a Δ*csrA* mutant on TY and M9/succinate agar and in TY and M9/succinate broth. (B) Desiccation tolerance of WT, Δ*csrA* mutant, Δ*csrA* mutant with pMMB (empty vector), and Δ*csrA* mutant with *csrA* expressed in trans (pAB2750) at day 0 and at day 6 of desiccation at 2% RH or at 6 days in PBS. The cell number represents the total number of viable cells recovered from each membrane. The data are the average of three or more biological replicates, and standard deviations are shown as error bars.

When desiccated after growth in M9/succinate to stationary phase, the AB5075 *ΔcsrA* mutant lost almost all viability over 6 days (Fig. 2B and Table 1). The desiccation phenotype was complemented by expressing *csrA in trans*. Δ*csrA* mutant cells incubated for 6 days after being filtered and resuspended in PBS remained fully viable (Fig 2B).

**Table 1.** Genes that contribute to desiccation tolerance of *A. baumannii* AB5075.

| Gene | Gene name | Gene annotation | Relative viability loss ratio (day 6/day 0 desiccation)<sup>a</sup> | Protein abundance (ratio of WT/Δ*csrA*) |
| --- | --- | --- | --- | --- |
| AB5075 (WT) |  |  | 1.0 |  |
| ABUW_0916 |  | Biofilm-associated protein | 1.5 | 2.6 |
| ABUW_2433 |  | KGG domain-containing protein | 125 | 4.3 |
| ABUW_2436 | *katE* | Catalase | 7.5 | 3.2 |
| ABUW_2437 |  | Heme oxygenase-like protein | 150 | 10.5 |
| ABUW_2639 |  | Universal stress protein family | 6.0 | 4.4 |
| ABUW_2724 |  | Hypothetical protein | 4.0 | 2.4 |
| ABUW_2750 | *csrA* | Carbon storage regulator | 480 | 8.5 |
| ABUW_3123 | *otsA* | Trehalose-6-phosphate synthase | 2.0 | 10.5 |
| ABUW_3346 | *acnA* | Aconitate hydratase 1 | 1.5 | 3.2 |

<sup>a</sup> All mutants were except the $\Delta$ *csrA* strain were grown in TSB broth to the stationary phase of growth prior to being desiccated. The viability loss of wt day6/day 0 was 17 for the WT grown in TSB broth. The viability loss of WT grown in M9/succinate was 276

### CsrA affects mutiple cellular processes in *A. baumannii*

To identify genes whose translation might be regulated by CsrA, we compared the proteomes of wild-type and Δ*csrA* cells (Table S2A). There were 97 proteins present at higher levels in the Δ*csrA* mutant compared to the wild type (ratio of Δ*csrA/WT* ≥2.5, Table S2B). Among these were proteins for type IV pilus assembly, synthesis of the siderophore ferric acinetobactin, and a glutamate/aspartate transporter. The Δ*csrA* mutant also had elevated levels of enzymes for the catabolism of hydroxcinnamates, phenylacetate and quinate. Levels of an alcohol dehydrogenase (ABUW_1621) and an aldehyde dehydrogenase (ABUW_1624) were also elevated. The Δ*csrA* mutant was defective in pilus-mediated twiching motility as assessed by movement aross a soft-agar plate (Fig. 3A). The mutant also had a severe growth defect when grown on succinate in the presence of ethanol (Fig. 3B). One possible explanation for this is that the *csrA* mutant metabolized ethanol to form toxic acetaldehyde to levels that slowed growth.

**Figure 3.**
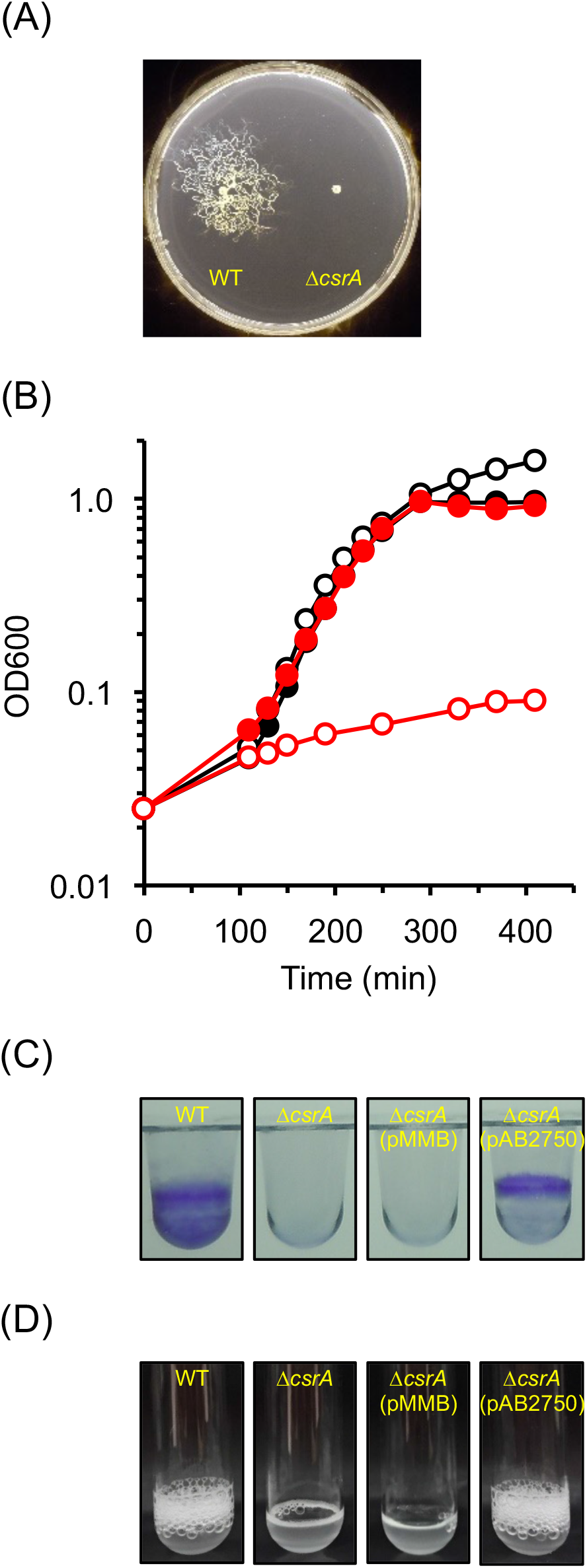
Phenotypes of the Δ*csrA* mutant: (A) twitching motility; (B) growth of wild type (black symbol) or the Δ*csrA* mutant (red symbol) in M9/succinate in the absence (closed symbol) or presence (open symbol) of 0.5% ethanol. The data shown are representative of each strain and condition; (C) crystal violet staining of biofilms from wild type, Δ*csrA* mutant, Δ*csrA* mutant with pMMB (empty vector), or Δ*csrA* mutant with pAB2750; and (D) catalase activity of wild type, Δ*csrA* mutant, Δ*csrA* mutant with pMMB (empty vector), or Δ*csrA* mutant with pAB2750.

There were 106 proteins present in lower amounts in the Δ*csrA* mutant compared to the wild type (ratio of WT/Δ*csrA* ≥ 2.5, Table S2C). A large proportion of these (39%) are annotated as hypothetical proteins. Several membrane proteins, and proteins annotated as involved in *β*-lactam antibiotic resistance (ABUW_1194, 2619, and 3497), trehalose synthesis (ABUW_3123) and possibly biofilm formation (ABUW_0916) were in lower abundance in the Δ*csrA* mutant compared to wild type. As reported previously, a Δ*csrA* mutant did not form biofilms (21). This phenotype was complemented by expressing the *csrA* gene *in trans* (Fig. 3C). The Δ*csrA* proteome profile also suggested that CsrA is invovled in promoting the expression of proteins invovled in oxidative stress, including peroxidase (ABUW_0628) and catalase (*katE*, ABUW_2436). Indeed the Δ*csrA* mutant lacked detectable catalase activity (Fig. 3D).

### Genes important for desiccation tolerance in *A. baumannii* AB5075

We took advantabge of the three-allele transposon library to test how important some of the gene transcripts that were likely to be controlled by CsrA were for desiccation tolerance (Tables 1 and S1). When possible, we tested two different transposon mutants (transposon insertions in different positions of the gene) for each gene. *A. baumannii* AB5075 produces opaque and translucent colony variants that interconvert at high frequency and reflect changes in the thickness of capsular exopolysaccharide (26). AB5075 cells with decreased capsule production are about 2.5-fold more sensitive to desiccation (27). Here, we used only opaque colonies of AB5075 and its mutant derivatives in our desiccation assays.

We found that *katE* and *ABUW_2639* mutants were about 5-fold more sensitive to desiccation than the wild type, whereas *ABUW_2433* and *ABUW_2437* mutants were greater than 100-fold more sensitive to desiccation than the wild type (Table 1, Fig 4). The phentoytpes of *ABUW_2433* and *ABUW_2437* mutants could be complemented (Fig S2). The ABUW_2433 protein has 411 amino acids and is annotated as a KGG domain-containing protein. The KGG domain comprises a small region in the N-terminus of the protein and the remainder of the protein is annotated by InterPro as a disordered region that includes a series of AT_hook DNA binding motifs (SMART SM00384). The full length ABUW_2433 sequence was predicted to be intrinsically unstructured when queried with the IUPred3 tool (https://iupred.elte.hu) (28). ABUW_2437 is annotated as an iron-containing redox enzyme or a heme-oxygenase-like protein (Fig 4). The predicted *ABUW_2437* transcript has traits characterisitic of a target of CsrA post-transcriptional regulation. The DNA sequence predicts a relatively long 5’ untranslated region (316 bp) and there is a predicted CsrA binding motif (GGA) in the ribosome binding site of the transcript. *ABUW_2639* is annoated as belonging to a universal stress protein A family. It has been shown to protect *A. baumannii* ATCC17987 from oxidative stress of hydrogen peroxide (29).

**Figure 4.**
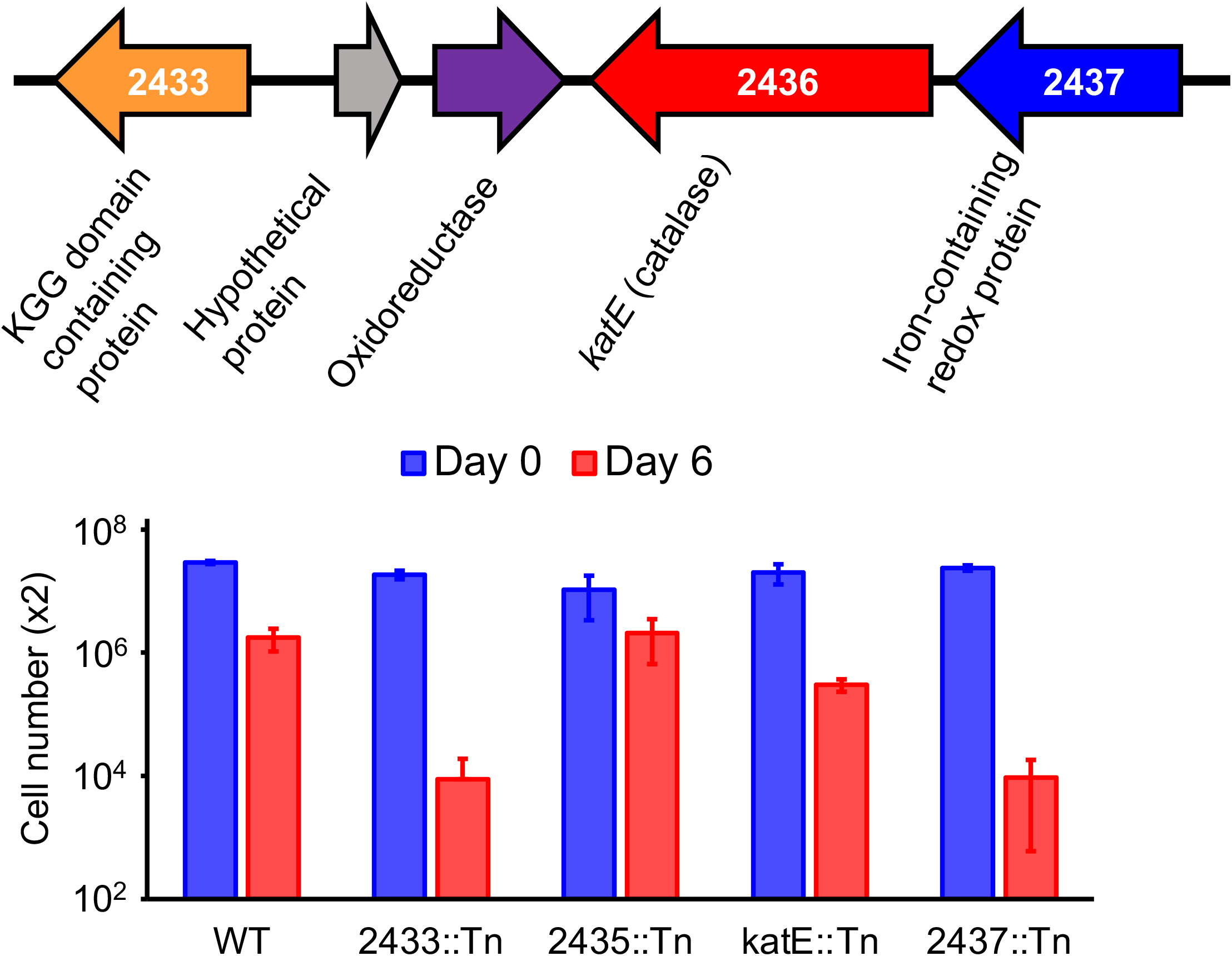
*A. baumannii* genes important for desiccation tolerance. (A) Genes and (B) Desiccation tolerance of mutants at 2% RH. The cell number represents the total number of viable cells recovered from each membrane. The data are the average of three or more biological replicates, and standard deviations are shown as error bars.

We wondered if *ABUW_2433* and *ABUW_2437* might play a role in promoting desiccation tolerance of the two *A. baumannii* strains, ATCC17978 and ATCC19606, that do not survive well at 2% RH (Fig 1B). ATCC19606 has the gene region shown in Fig 4 intact, but the gene that is homologous to *ABUW_2433*, encoding the KGG domain-containing protein, is annotated as a pseudogene. ATCC17978 appears to be missing a gene in homologus to *ABUW_2433*. However it has conitguous *katE* and iron-containing redox protein genes (*A1S_1386 and A1S_1385*). Expression of the two AB5075 genes *in trans* improved the survival of the two ATCC strains at 2% RH (Fig 5), providiing evidence that ABUW_2433 and ABUW_2437 are generally important for desiccaton tolerance.

**Figure 5.**
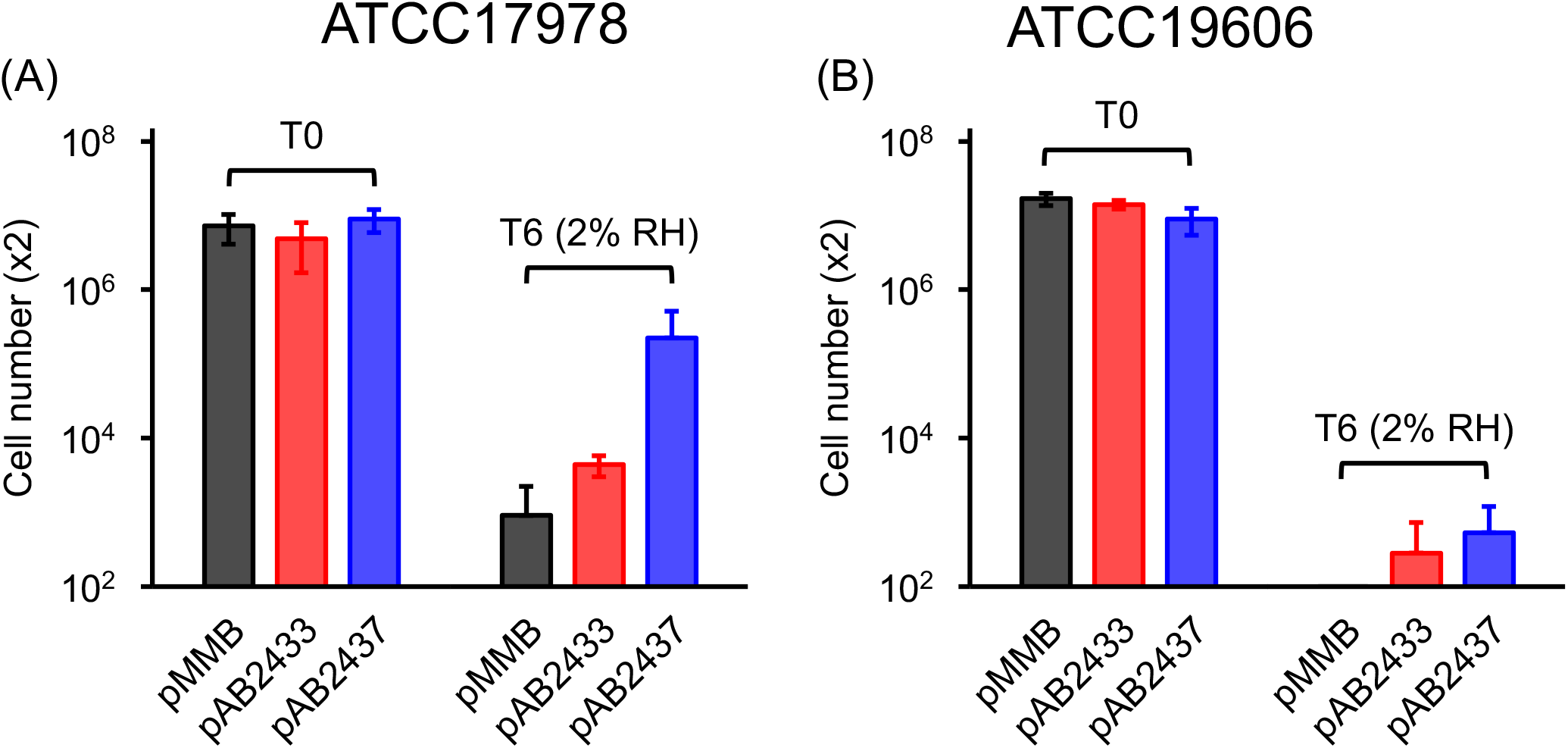
Effect of expression of *ABUW_2433* (red bars) or *ABUW_2337* (blue bars) gene *in trans* on desiccation tolerance of *A. baumannii* (A) ATCC17978 and (B) ATCC19606 after 0 days (control) and 6 days of desiccation at 2% RH. Empty vector (pMMB, black bars) was used as a control. The data are the average of three or more biological replicates, and standard deviations are shown as error bars.

### Other possible desiccation tolerance genes

As shown in Table 1, we identified an additional six genes that are likely regulated by CsrA, that may have a small role in desiccation tolerance. *ABUW_0916*, encoding a biofilm-associated protein and *otsA*, encoding tehalose-6-phosphate synthase, are the only two of the six for which we can hypothesize some connection to desiccation. Biofilms have been shown to be important for desiccation tolerance of bacteria (15, 16) and trehalose has been shown to play a significant role in desiccation tolerance of eukaryotes and bacteria (30, 31). When added extrinsically to cultures, trehalose increased the desiccation tolerance of *A. baumannii* ATCC 19606 (22). However, a *ΔmtlD-otsB* mutant of ATCC19606, defective in endogenous production of the compatible solutes, mannitol, and trehalose, was not more sensitive to desiccation than the wild type (22).

## DISCUSSION

Depletion of water during desiccation leads to loss of membrane integrity and accompanying disruption of aerobic respiration results in the generation of reactive oxygen species, including hydrogen peroxide (32). So, it makes sense that *katE*, encoding catalase, contributes to desiccation tolerance. Proteomics analyses of *A. baumannii* showed that proteins involved in redox defense including catalase, alkyl peroxidase reductases and superoxide dismutase were elevated in stationary-phase cells (33), which is consistent with the observation made by many that cells stationary-phase cells survive desiccation much better than exponentially growing cells.

Since the desiccation-tolerance genes *ABUW_2433* and *ABUW_2437* are near or adjacent to *katE*, on the genome and all are likely to be controlled by CsrA, it seemed important to consider that they might somehow mediate oxidative stress tolerance even though the amino acid sequences of the encoded proteins don’t have motifs typically associated with reactive oxygen species detoxification. However, we were unable to demonstrate that *ABUW_2433*::Tn and *ABUW_2437*::Tn mutants were sensitive to hydrogen peroxide and paraquat - both powerful oxidizing agents. In addition, a study that looked at effects of hydrogen peroxide exposure on gene expression in *A. baumannii*, found that *katE* but not *ABUW_2433* or *ABUW_2437*, was expressed at elevated levels and neither of these genes is part of the OxyR regulon that controls the response to oxidative stress in *A. baumannii* (34). Thus, we do not think that either of these proteins, which have the greatest defects in desiccation found to date, are likely to function by protecting cells against oxidative stress

The predicted physical properties of ABUW_2433 provide suggestions as to how it may function. It is an intrinsically disordered protein (IDP) that is highly hydrophilic, with 27% positively charged amino acids residues and 31% negatively charged residues. It is also predicted to assume a collapsed or extended conformation, likely depending on its context (ROBETTA PFRMAT TS prediction; https://robetta.bakerlab.org). ABUW_2433 has 13 repeated AT-hook DNA binding motifs that extend across about 70% of the protein. This motif preferentially binds to AT-rich sequences in the minor groove of DNA. AT-hook DNA binding motifs are found primarily in eukaryotic proteins, many of which have roles in transcriptional regulation (35, 36). Only 8.5% of annotated AT hook DNA binding motifs are found in bacteria, but about half of these are found in gamma proteobacteria, the group to which *A. baumannii* belongs. We hypothesize that ABUW_2433 binds to *A. baumannii* DNA and somehow protects it from desiccation-induced damage. IDPs are critical for the microscopic animals called tardigrades to survive desiccation. When desiccated, some of these proteins vitrify and probably trap desiccation sensitive molecules in a noncrystalline amorphous matrix, thereby protecting them from denaturation or other forms of destruction (37, 38). IDPs or proteins with IDP domains are less common in prokaryotes than in eukaryotes, but drawing from work on eukaryotes, they are proposed to play a central role in cellular process in bacteria that may depend on the formation of molecular condensates (39). It is possible that this is important for the viability of desiccated *A. baumannii*.

Although not much work has been done on *Acinetobacter* CsrA, based on what is known for other gamma proteobacteria, we hypothesize that a set of ncRNAs that is induced by a GacSA (ABUW_3306 and ABUW_3639) two-component regulatory system, controls the repressor activity of CsrA by sequestering it (40). We can draw a link between the GasSA system and CsrA because they both appear to control catabolism of the aromatic compound phenylacetate. An *A. baumannii* Δ*gacA* mutant is unable to catabolize phenylacetate (41), and our proteomics results suggest that CsrA acts to repress the synthesis of at least one enzyme required for phenylacetate degradation. In fact, we have determined that the Δ*csrA* mutant grows better on phenylacetate; with a doubling time of 39 min, than the wild type, which has doubling time of 50 min. We hypothesize that a Δ*gacA* mutant does not synthesize ncRNAs that would normally “sponge-up” CsrA, thus allowing CsrA to bind to the 13-gene (ABUW_2524 to ABUW_2536) phenylacetate mRNA transcript to repress its translation. At this point we do not have a clear understanding of the inventory of *A. baumannii* ncRNAs that may bind to CsrA, but ncRNAs are abundant in AB5075, and several of them are expressed at extremely high levels (42). The desiccation phenotype of CsrA appears to depend on its ability to activate translation and although it’s difficult to reconcile this activity with a model where CsrA is sequestered by ncRNAs, it is known that ncRNA turnover can occur resulting in the release of free CsrA (43). Most of what is known about mechanisms of CsrA action centers on its role as a repressor of translation (44–47) and it may be of interest to probe its capability as an activator in *A. baumannii*.

We found that *A. baumannii* AB5075 survived desiccation for six days at 2% RH much better than two other *A. baumannii* strains that we tested, but it is important to note that most studies of desiccation tolerance have been carried out at about 30% RH or in room air and the emphasis has been on the number of days or months that a particular strain remains viable when desiccated. When Farrow et al (20) tested the survival of several strains that were dried and incubated at an RH of 25–61% (mean 46%) they found AB5075 to have an average survival time of 90 days, whereas strains ATCC19606 and ATCC17978 had average survival times of 3 and 34 days respectively. Even though AB5075 is tolerant to desiccation over months at a mean RH of 46% and over days at 2% RH, we cannot necessarily conclude that the same sets of genes are needed for desiccation tolerance under these two different conditions. For example, Farrow et al. (20) found that the response-regulator protein BmfR was important for desiccation tolerance of ATCC17978 in long term desiccation assays, whereas we did not observe a role of *bmfR* in protecting AB5075 from desiccation in shorter term incubations at 2%RH (Table S1).

Here we established a robust assay for desiccation tolerance of a highly virulent strain of *A. baumannii* and identified two genes, *ABUW_2433* and *ABUW_2437*, that are extremely important for desiccation tolerance. Outside of the *Acinetobacter* genus, *ABUW_2437* has homologs in *Pseudomonas stutzeri* (40% amino acid identity). However, except for a partial homolog found in *Enterobacteriaceae* bacterium TzEc051 (99% identity over 38% of the AB5075 protein), *ABUW_2433* does not appear outside *Acinetobacter*. This and the unusual predicted physical properties of ABUW_2433 as anIDP, suggest that there is novel physiological basis for desiccation tolerance in *A. baumannii* that remains to be explored.

## MATERIALS AND METHODS

### Bacterial strains and growth conditions

Strains used in this study are listed in Table S3A. Strain AB5075 (AB5075-UW) was used as a wild type (24) and individual transposon mutants were obtained from the Manoil lab comprehensive ordered transposon mutant library at the University of Washington (24). All strains except for the Δ*csrA* mutant were routinely grown and maintained in TY (10 g Tryptone, 5 g Yeast extract, and 8 g NaCl in 1000 ml) medium or BBL Trypticase Soy Broth (TSB) media at 37°C, unless otherwise stated. The Δ*csrA* mutant was grown in M9/succinate.

### Desiccation assay

Strains from a frozen stock (−80°C) were streaked onto TY plates and incubated at 37°C. Colonies (three to five) were picked and inoculated into 2 ml of TSB, and cultures were grown overnight at 37°C with a shaking speed of 200rpm. Overnight cultures were diluted to yield an initial OD_600_ of 0.025 in 10 ml TSB in a 50 ml Erlenmeyer flask. Cultures were grown at 37°C with a shaking speed of 200 rpm to mid-exponential-phase (OD_600_=0.4 to 0.6) or stationary-phase (24 hours after inoculation). Cells were harvested by centrifugation and washed twice with Dulbecco’s phosphate-buffered saline (DPBS, Gibco), and cell density was adjusted to OD_600_=1 (about 5 x 10^8^ cells/ml) with DPBS. Cell suspension (2 spots of 50 μl each per membrane) was filtered onto a 0.4 μm Whatman nucleopore polycarbonate track-etched membranes (25 mm diameter) that had been placed in Nalgene analytical filter units, and the membranes were then placed into 15 ml uncapped centrifuge tubes. To obtain the T0 (baseline) viable cell number, 1 ml of DPBS was immediately added to one tube and incubated for 5 min at room temperature (24 ± 2°C) on a rotary shaker. Viable cell numbers were determined by plating on TSB agar. For desiccation, tubes with membranes were placed in a Snapware containers (2.3 x 6.3 x 8.4 inches) that contained DRIERITE in the lids of 50 ml centrifuge tubes (x4, 7.5 g of DRIERITE desiccant in each lid) or saturated calcium chloride hexahydrate solution in 5 ml beaker (x8) to yield the RHs of 2% or 30% (± 2), respectively. The Snapware containers were incubated at room temperature. Digital hygrometers (VWR International Ltd) were placed in each container to monitor the RH. At desired time points, tubes containing membranes were removed from the containers, 1 ml of DPBS was added to each tube, and incubated for 5 min at room temperature on a rotary shaker. Viable cell counts were determined on TSB agar. For each strain, a minimum of three biological replicates of desiccation assays were performed except for individual transposon mutants, which were assayed twice for each allele.

### Construction of the Δ*csrA* mutant

In-frame deletion of the *csrA* (*ABUW_2750*) gene was generated by overlap extension PCR as described (48). PCR primers are listed in Table S3B. PCR product was cloned into mobilizable suicide vector pEX2-TetRA and transformed into *E. coli* NEB 10-beta (New England Bio Labs). The sequence-verified deletion construct was transformed into *E. coli* S17-1, and further mobilized into *A. baumannii* strain AB5075 by conjugation on TY agar. Single recombinant conjugants were first selected on M9/succinate plate containing 20 μg/ml Tc, and Tc resistant colonies were further plated onto M9/succinate plate containing 5% sucrose. Sucrose resistant and Tc sensitive colonies were screened by colony PCR and sequencing to validate the expected chromosomal in-frame deletion of the *csrA* gene.

To complement the Δ*csrA* mutant, the full length *csrA* gene plus the 15 bp upstream that contains the putative ribosome binding site was PCR amplified and cloned into pMMB67EH-TetRA. The construct was transformed into *E. coli* NEB 10-beta (New England Bio Labs). The sequence-verified construct was transformed into *E. coli* S17-1, and further mobilized into the Δ*csrA* mutant by conjugation on M9/succinate plates containing 20 μg/ml Tc. As a negative control, empty vector pMMB67EH-TetRA was used. The same procedure was used to clone *ABUW_2433* and *ABUW_2437* for complementation experiments.

### Phenotypic characterization of the Δ*csrA* mutant

M9/succinate was used in all experiments. For motility assays, an overnight culture (16 to 18 hours) was diluted to yield OD_600_=0.5, and 2 μl of sample was spotted onto the freshly prepared M9/succinate plate and incubated at 37°C for 24 h. For biofilm assays, an overnight culture was inoculated into 100 μl of M9/succinate in Costar vinyl 96 well “U” bottom plates (initial OD_600_=0.05), and the plates were sealed with Breath-Easy sealing membranes. After incubation at room temperature for 48 h, culture was removed, the plate was rinsed with tap water twice, and 150 μl of 0.1% crystal violet solution was added to each well. After incubating at room temperature for 15 min, crystal violet solution was removed, the plate was rinsed with tap water 5 times, and the plate was dried at room temperature. For catalase assays, cells were harvested at OD_600_=0.5, supernatant was removed, and cells were resuspended in DPBS to yield 10 mg wet cell/100 μl DPBS. 100 μl of cell suspensions were placed in 13 x 100 mm borosilicate glass tubes. Then 100 μl of 1% Triton X-100 and 100 μ1 of 30% hydrogen peroxide were added, mixed thoroughly, and incubated for 15 min at room temperature (49).

### Label-free protein quantification

Since the Δ*csrA* mutant had a severe growth defect on TY medium, both the wild type and Δ*csrA* mutant were grown in M9/succinate. Two biological sample replicates were prepared for each strain. Cells from each culture were harvested at OD_600_=0.5 by centrifugation, washed twice with DPBS, and cells were stored in −80°C before further analysis. Cells were lysed in buffer containing 4% SDS, 100 mM Tris pH8.0, 10 mM DTT by heating at 95°C for 5 min. After cooling to room temperature, the lysates were sonicated with ultrasonication probe on ice to shear DNA. Total protein concentration was determined by the BCA assay (Thermo Pierce, Rockford, IL). 500 μg of each protein lysate was reduced and alkylated, diluted in 8 M urea solution, and the SDS was removed with a 3kD molecular weight cutoff filter. After buffer exchange, the protein lysates were digested with trypsin (Promega, Madison, WI) at 37°C overnight and the digested samples were desalted with 1cc C18 Sep-Pak solid phase extraction cartridges (Milford, MA, Waters). The eluted samples were vacuum dried and resuspended in 0.1% formic acid. Reverse phase nanoLC-MS analysis of the protein digests was carried out with a Thermo Easy-nanoLC coupled to a Thermo Q-Exactive Plus Orbitrap mass spectrometer. Triplicate top 20 data-dependent acquisition runs were acquired for each sample, and 1 μg of protein digest was loaded for each run. The peptides were separated by a 50 cm x 75 μm I.D C8 column (5 μm diameter, 100 Å pore size C8 MichromMagic beads) with a 90 min 10 to 30% B gradient (solvent A: 0.1% formic acid in water, solvent B: 0.1% formic acid in acetonitrile, flow rate 300 nl/min). The MS data acquisition parameters were set as follows: full MS scan resolution 70k, maximum ion injection time 100 mS, AGC target 10^6^, scan range of 400 to 2000 m/z; MS/MS scan resolution 17.5 k, maximum ion injection time 100 mS, AGC target 5^4^, isolation window 1.6 m/z, HCD NCE 35 scan range of 200 to 2000 m/z; loop count 20, intensity threshold 5^3^, underfill ratio 1%, dynamic exclusion 10 sec. High resolution MS^2^ spectra were searched against a target-decoy proteome database of strain AB5075 (a total of 7678 sequences) downloaded from Uniprot (Oct17, 2017) using Comet (version 2015.02 rev. 1) (50) with following parameters: precursor peptide mass tolerance 20 ppm, allowing for −1, 0, +1, +2, or +3 ^13^C offsets; fragment ion mass tolerance 0.02 Da; static modification, carbamidomethylation of cysteine (57.0215 Da); variable modification, methionine oxidation (15.9949 Da). The search results were further processed by PeptideProphet (51) for probability assignment to each peptide-spectrum match, and ProteinProphet (52) for protein inference and protein probability modeling. The output pepXML files from three technical replicates were grouped for subsequent spectral counting analysis using Abacus (53). The pepXML and protXML files for each sample, combined ProteinProphet file from all samples were parsed into Abacus for spectral counting of each protein. The following filters were applied for extracting spectral counts from MS/MS datasets: (1) the minimum PeptideProphet score the best peptide match of a protein must have maxIniProbTH=0.99; (2) The minimum PeptideProphet score a peptide must have to be even considered by Abacus, iniProbTH=0.50; (3) The minimum ProteinProphet score a protein group must have in the COMBINED file, minCombinedFilePw=0.90. Spectral counts for 1616 proteins were reported across four sample groups (two strains and two biological replicates) with estimated protein false discovery rate of 1.94%. The protein expression fold changes between wild type AB5075 and Δ*csrA* mutant were computed from adjusted spectral counts output from Abacus. The mass spectometry data have been deposited to the ProteomeXchange Consortium via the PRIDE partner repository with the dataset identifier xxxxxxxxx..

## ACKNOWLEDGMENTS

We thank Dr. Hemantha Don Kulasekara and Dr. Samuel Miller (University of Washington) for sharing the pEX2-TetRA and pMMB67EH-TetRA vectors. We thank Indranil Biswas for alerting us to the possibility that intrinsically disordered proteins could be involved in desiccation tolerance in *A. baumannii*.

## FUNDING

This work was supported by the Functional Genomics Program, National Institute of Allergy and Infectious Diseases under Grant 1U19AI107775-01.

## SUPPLEMENTAL MATERIAL

**Figure S1.**
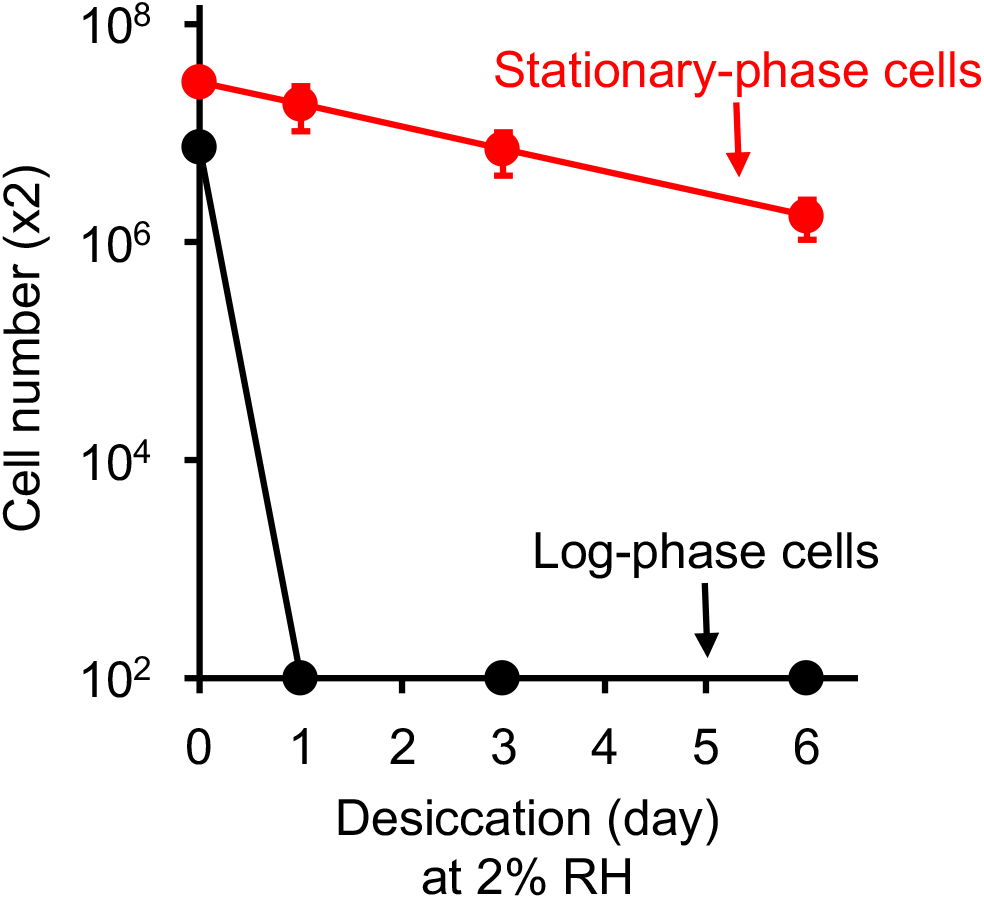
Desiccation tolerance of *A. baumannii* AB5075 cells from the log- and stationary-phases of growth. The cell number represents the total number of viable cells recovered from each membrane. The data are the averages of three or more biological replicates, and standard deviations are shown as error bars.

**Figure S2.**
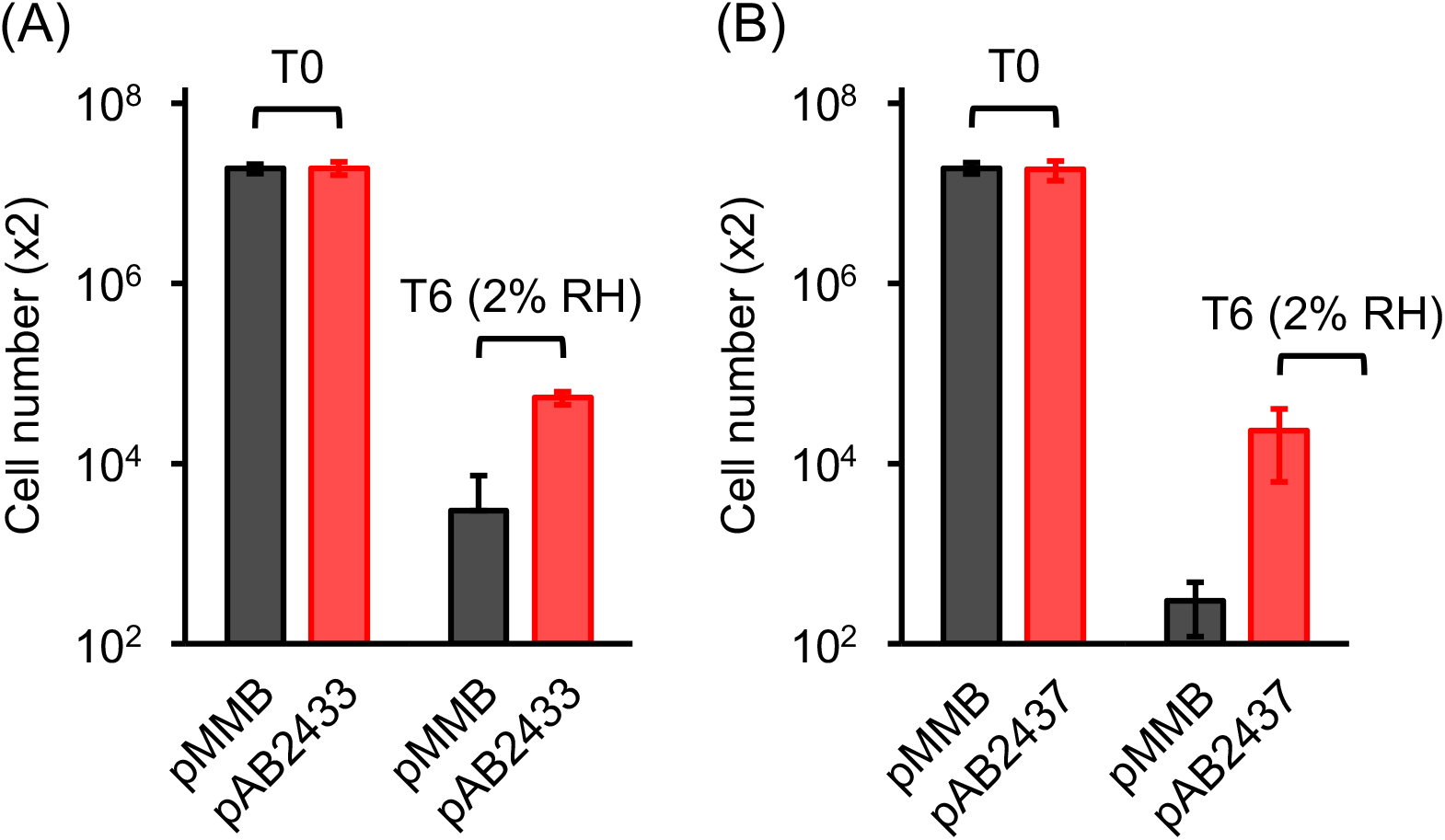
Complementation of desiccation phenotypes of (A) *ABUW_2433*::Tn and (B) *ABUW_2437*::Tn after 0 days (control) and 6 days of desiccation at 2% RH. Empty vector (pMMB) was used as a control. The data are the average of three or more biological replicates, and standard deviations are shown as error bars.

**Table S1.** Mutants from the three-allele library tested for desiccation tolerance

**Table S2.** (A) Label-free protein quantification of wild type and the Δ*csrA* mutant. (B) List of proteins up-regulated (ratio of Δ*csrA*/WT ≥2.5 and Δ*csrA* read count ≥7.5) in the Δ*csrA* mutant compared to wild type. (C) List of proteins down-regulated (ratio of Δ*csrA*/WT ≤0.4 and WT read count ≥7.5) in the Δ*csrA* mutant compared to wild type.

**Table S3.** (A) Bacterial strains used in this study. (B) Plasmids and primers used in this study

## Notes

### Competing Interest Statement

The authors have declared no competing interest.

### Summary of Updates

no data have been changed, but some data have been removed and the MS has been rewritten for clarity.

